# Male and female mice demonstrate divergent cellular responses in the bed nucleus of stria terminalis (BNST) following morphine withdrawal

**DOI:** 10.1101/369173

**Authors:** Brennon R. Luster, Elizabeth S. Cogan, Karl T. Schmidt, Dipanwita Pati, Melanie M. Pina, Kedar Dange, Zoé A. McElligott

## Abstract

The United States is experiencing an opioid epidemic of significant proportions, imposing enormous fiscal and societal costs. While prescription opioid analgesics are essential for treating pain, the cessation of these drugs can induce a withdrawal syndrome, and thus opioid use often persists to alleviate or avoid these symptoms. Therefore, it is essential to understand the neurobiology underlying this critical window of withdrawal from opioid analgesics to prevent continued usage. To model this, we administered a low dose of morphine, and precipitated withdrawal with naloxone to investigate the behavioral and cellular responses in C57BL/6J male and female mice. Following 3 days of administration, both male and female mice sensitized to the repeated bouts of withdrawal, as evidenced by their composite global withdrawal score. Female mice exhibited increased withdrawal symptoms on some individual measures, but did not show characteristic weight loss observed in male mice. Because of its role in mediating withdrawal-associated behaviors, we examined neuronal excitability and inhibitory synaptic transmission in the bed nucleus of the stria terminalis (BNST) 24 hours following the final precipitated withdrawal. In male mice, morphine withdrawal increased spontaneous GABAergic signaling compared to controls. In contrast, morphine withdrawal decreased spontaneous GABAergic signaling, and increased BNST projection neuron excitability in female mice. Intriguingly, these opposing GABAergic effects were dependent on within slice excitability. Our findings suggest that male and female mice manifest divergent cellular responses in the BNST following morphine withdrawal, and alterations in BNST inhibitory signaling may be a significant factor contributing to the expression of behaviors following opioid withdrawal.

## Introduction

The United States is currently experiencing an opioid epidemic that claims the lives of an estimated 115 Americans every day (Center for Disease Control, 2017). Drug overdoses continue to rise in the U.S., and more than three out of five drug overdose deaths involve an opioid (Hedegaard et al., 2016). Prescription opioids are useful in treating acute pain, however, the cessation of these drugs, especially after chronic use, can induce a highly dysphoric withdrawal syndrome marked by negative somatic (vomiting, cold/hot flashes, shaking, diarrhea, etc.) and emotional symptoms (anxiety, dread) which often lead patients into persistent opioid seeking. Indeed, intensity of withdrawal symptoms is highly correlated to poor patient compliance for opioid use disorder (OUD) (Fishman, 2008; Kanof et al., 1993), and increased Distress Intolerance (the actual and/or perceived lack of ability to cope with the aversive somatic and/or emotional states associated with withdrawal) is associated with elevated opioid misuse in chronic pain patients (McHugh et al., 2016). Additionally, prescription opioid exposure alone is a strong risk factor for an OUD. It has been shown that patients with chronic noncancer pain who were prescribed opioid analgesics had significantly higher rates of developing an OUD compared to those not prescribed opioids (Edlund et al., 2014). Therefore, it is imperative to investigate this critical period of withdrawal from prescription opioid analgesics which often initiates opioid misuse.

Furthermore, there are a growing number of studies that highlight the role of sex in opioid use and misuse. Studies show that women have a higher incidence of pain (Tsang et al., 2008) and report higher pain intensity compared with men (Arendt-Nielsen et al., 2004; Eriksen et al., 2003; Greenspan et al., 2007). Women are also more likely to be prescribed opioids and given higher doses for longer periods of time (Kelly et al., 2008; McCabe et al., 2005; Simoni-Wastila et al., 2004; Sullivan et al., 2005), whereas men demonstrate more substance abuse behavior than women (Kessler et al., 2005; Stinson et al., 2005). Women also report significantly higher cravings for opioids (Lee and Ho, 2013). Despite these findings, there is a lack of pre-clinical studies investigating sex as a biological variable in key brain nodes that are engaged during opioid exposure and withdrawal.

The bed nucleus of the stria terminalis (BNST), a component of the extended amygdala, is a ventral forebrain structure containing a large population of gamma-aminobutyric acid (GABA) neurons. The BNST functions as a critical relay center to regulate a range of emotional and motivational processes, receiving inputs from cortical and limbic structures, and via reciprocal projections with several limbic and hindbrain nuclei (Dong and Swanson, 2004, 2006; McElligott and Winder, 2009; Sun and Cassell, 1993). Recent studies have shown that GABAergic projection neurons from the BNST to the ventral tegmental area (VTA) and lateral hypothalamus (LH) drive reward-like behavior (Jennings et al., 2013a,b), and neuromodulation enhances fear and anxiety-like responses in the BNST by inhibiting these output neurons (Marcinkiewcz et al., 2016). The BNST plays a critical role in the negative valence associated with opioid withdrawal (Delfs et al., 2000) and inactivation of the BNST has been shown to block morphine withdrawal-potentiated startle (Harris et al., 2006). Moreover, the BNST is a highly sexually dimorphic brain region (Hisasue et al., 2010; Tsuneoka et al., 2017). These anatomically sexually dimorphic observations, however, were focused in the posterior division of the BNST where female mice contain roughly half the number of neurons as male mice (Shah et al., 2004; Xu et al., 2012), rather than the anterior portion of the BNST, which is thought to modulate drug seeking and motivational states. To date, there have not been detailed physiological studies investigating cellular responses to opioids in the anterior portion of the BNST in both male and female mice. Thus, the objective of this study is to examine sensitizing repeated bouts of withdrawal from an opioid analgesic, and to investigate the molecular modifications induced in the BNST of male and female mice.

## Materials and Methods

All experiments were performed in accordance with the University of North Carolina at Chapel Hill (UNC) Institutional Animal Care and Use Committee’s guidelines. C57BL/6J male and female mice (aged 7 weeks; Jackson Laboratories) were group or signally housed within UNC animal facilities on a normal light cycle (lights on at 7 am, lights off at 7 pm) and given food and water *ad libitum*.

### Morphine Withdrawal Paradigm

To model morphine withdrawal we used 4 groups of animals: morphine-naloxone males (MNM), saline-naloxone males (SNM), morphine-naloxone females (MNF), and saline-naloxone females (SNF). To model withdrawal from opioid analgesics, we administered a naloxone-precipitated morphine withdrawal paradigm (Fig. 1A). Mice were injected once daily at approximately 9am with morphine (10 mg/kg) or saline (0.1 mL/g) subcutaneously (s.c.), followed 2 h later by naloxone (1 mg/kg) s.c. These injections were repeated for 3 days. Following naloxone administration, mice were observed for 10 min and the following individual withdrawal symptoms were tallied and recorded: escape jumps, teeth chattering, wet dog shakes, paw tremors, diarrhea, excessive eye blinking, genital grooming, ptosis, and abnormal posturing. A daily composite global withdrawal score was determined for each mouse by adding the total number of escape jumps to all other withdrawal symptoms which were scored as either a 1 (present) or 0 (absent) (McNally and Akil, 2002).

**Figure 1:**
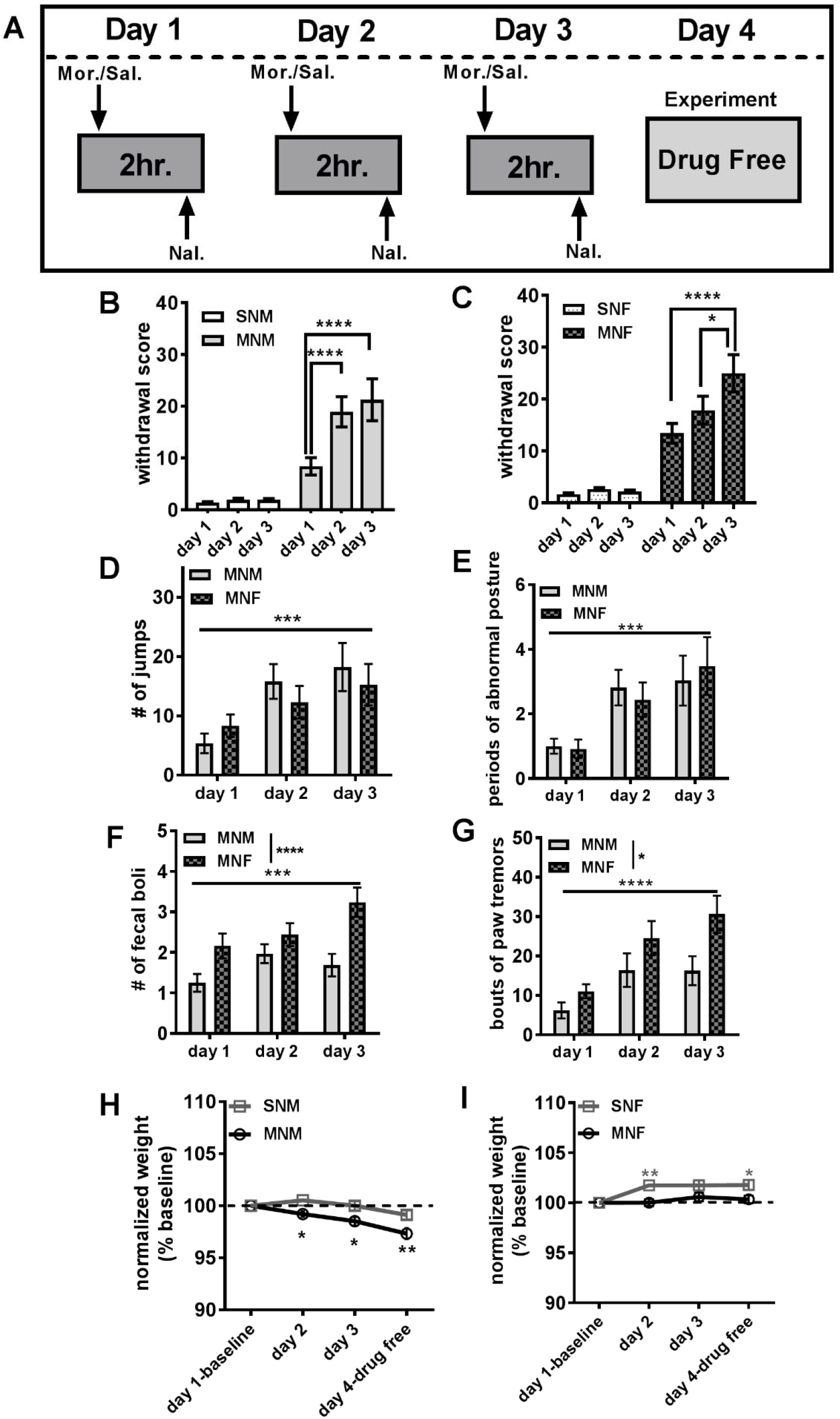
Morphine withdrawal in male and female mice **A)** Naloxone-precipitated morphine withdrawal paradigm. **B)** Average withdrawal scores in male mice (SNM n=22; MNM n=18). **C)** Average withdrawal scores in female mice (SNF n=15; MNF n=15). **D)** The average number of jumps between MNM and MNF. **E)** The average number of fecal boli produced between MNM and MNF. **F)** The average number of paw tremors between MNM and MNF. **F)** The average number of paw tremors between MNM and MNF. **G)** The average number of wet dog shakes between MNM and MNF. **H)** Average weight loss in male mice (% baseline). **I)** Average weight loss in female mice (% baseline). Each bar represents the mean ± SEM, * (p<0.05), ** (p<0.01), *** (p<0.001), **** (p<0.0001).

### Brain slice preparation whole-cell electrophysiology

All electrophysiology experiments were performed approximately 24 h following the last naloxone administration. Mice were anesthetized (isoflurane) and decapitated as previously described (McElligott et al., 2010). Briefly, brains were quickly removed and placed in ice-cold (1-4° C) high-sucrose artificial cerebral spinal fluid (ACSF) (in mM: 194 sucrose, 20 NaCl, 4.4 KCl, 2 CaCl_2_, 1 MgCl_2_, 1.2 NaH_2_PO_4_, 10 glucose, 26 NaHCO_3_) that had been oxygenated (95% O_2_, 5% CO_2_) for a minimum of 15 minutes. Coronal slices (300 μm) containing the BNST were prepared using a Leica VT1000 (Leica, Germany), and slices were allowed to equilibrate in oxygenated ACSF (in mM: 124 NaCl, 4.4 KCl, 2 CaCl_2_, 1.2 MgSO_4_, 1 NaH_2_PO_4_, 10 glucose, 26 NaHCO_3_, 34° C) for at least 1 hour. Slices were then transferred to the submerged recording chamber and perfused with oxygenated ACSF (28-30° C) at a rate of 2 ml/min.

### Whole-Cell Recordings

Slices were placed in a submerged chamber (Warner Instruments). BNST neurons were directly visualized with an upright video camera (Hamamatsu). Recording electrodes (3–6 MΩ) were pulled on a P-97 Flaming-Brown Micropipette Puller (Sutter Instruments) using thin-walled borosilicate glass capillaries. Spontaneous or miniature GABA_A_-mediated inhibitory postsynaptic currents (sIPSCs or mIPSCs) were acquired in voltage-clamp at −80mV holding potential in 5-minute blocks (Kash and Winder, 2006). Recording electrodes to examine IPSCs were filled with cesium chloride (CsCl) intracellular recording solution (in mM: 130 CsCl, 1 EGTA, 10 HEPES, 2 ATP, 0.2 GTP) in the presence of kynurenic acid (3mM), to block ionotropic glutamatergic receptor currents. mIPSCs were recorded in the presence of kynurenic acid and tetrodotoxin (TTX, 500nM) to block activity dependent synaptic release. Electrically evoked-IPSCs (eIPSCs) were measured by local stimulation (0.1ms duration, two pulses) using a bipolar nichrome electrode while neurons were voltage-clamped at −70 mV. Stimulation intensity was adjusted to obtain approximately 80% of the maximum response. Recordings examining neuronal excitability were performed in current-clamp using potassium gluconate (KGluc) intracellular recording solution (in mM: 135 KGluc, NaCl 5, MgCl_2_ 2, HEPES 10, EGTA 0.6, Na_2_ATP 4, Na_2_GTP 0.4). Signals were acquired via a Multiclamp 700B amplifier (Molecular Devices), digitized and analyzed via pClamp 10.6 software (Molecular Devices). Input resistance, holding current, and access resistance were all monitored continuously throughout the duration of experiments. Experiments in which changes in access resistance were greater than 20% were not included in the data analysis.

The BNST contains both interneurons, and neurons the project to classical reward nuclei like the VTA and LH, which we term projection neurons (BNST_proj_). The projection neurons have characteristics of low capacitance, high input resistance, and lack I_h_ potassium channel current (Kash and Winder, 2006; Dumont and Williams, 2004 Marcinkiewcz et al., 2016). In order to investigate the effects of our morphine withdrawal paradigm on this circuitry, we analyzed cells that displayed these distinct physiological characteristics defined by the absence of Ih current, small membrane capacitance (average 32.2 ± 0.8 pF), and high input resistance (average 1060 ± 38 MΩ).

### Drugs

Morphine sulfate (non-selective opioid receptor agonist) and naloxone HCl (non-selective opioid receptor antagonist) were purchased from Sigma Aldrich (www.sigmaaldrich.com). Tetrodotoxin (TTX; sodium channel blocker) and kynurenic acid (non-selective glutamate receptor antagonist) were purchased from Abcam (www.abcam.com). Morphine sulfate and naloxone HCl were dissolved in sterile saline (0.9%) prior to injection. All drugs were injected at a volume of 0.1mL/10g.

### Data Analysis

Data are presented as mean ± SEM or cumulative distributions ± SEM. All statistical analyses were performed using GraphPad Prism version 7.03 for Windows (www.graphpad.com). A Student’s unpaired t-test or 2-way ANOVA was performed where appropriate. If a significant interaction was detected in the 2-way ANOVA, a post-hoc Tukey HSD or Holm-Sidak test was performed.

## Results

### Morphine withdrawal in males and females

To model repeated opioid withdrawal, mice were subjected to a morphine withdrawal paradigm (Fig. 1A) modified from studies performed in rats (McElligott et al., 2013; Schulteis et al., 1999). Mice were injected (s.c.) with either 10 mg/kg morphine (in 0.9% saline) or equal volume saline and returned to their homecage. 2 hours later all mice were injected (s.c.) with 1 mg/kg naloxone (in 0.9% saline) and placed in a clear 58.4 cm X 41.3 cm X 31.4 cm plastic bin. Following administration of naloxone, we observed mice for 10 min and a withdrawal score was calculated (Fig. 1B & 1C). In males, the global withdrawal score had a significant drug by day interaction (F(2,120)=8.054, p<0.001). MNM mice (n=32) demonstrated a sensitization of the global withdrawal score, with day 2 (p<0.0001) and day 3 (p<0.0001) significantly increased compared to day 1 (Fig. 1B). Similarly, data from MNF (n=25) showed a drug by day interaction (F(2,96)=4.751, p<0.05) with day 3 significantly increased compared to day 1 (p<0.0001), and day 3 significantly increased compared to day 2 (p<0.05). While SNM (n=30) and SNF (n=25) mice exhibited a few signs of withdrawal following naloxone administration, there was no sensitization of this withdrawal over time.

Because recent evidence has demonstrated disparities in behavioral responding between male and female rodents (Becker and Chartoff, 2018; Shansky, 2018), we next compared the individual withdrawal behaviors between our MNM and MNF mice across the 3 days of treatment to note any potential sex differences. Both MNM and MNF mice exhibited escape jumps and periods of abnormal posture, where there was a main effect of day (p<0.001 both measures; escape jumps: F(2, 110)=9.678, abnormal posture: F(2,110)=10.46; Fig. 1D and E) but not of sex (jumps: p=0.722; abnormal posture: p=0.998). In contrast, MNF mice produced significantly more fecal boli (main effect of sex F(1,165)=18.36, P<0.0001, Fig. 1E), and exhibited more paw tremors than MNM mice (main effect of sex F(1,162)=9.34, P<0.01, Fig.1F). While both male and female animals exhibited symptoms of teeth chattering, genital grooming, swallowing movements, and wet dog shakes, we did not observe differences in day of withdrawal or sex across these measures. We next evaluated changes in body weight following each bout of withdrawal and on the 4^th^ day prior to euthanasia. MNM mice exhibited on average over 2.5% loss in body weight across the days of treatment (Fig. 1H). There was a significant drug by day interaction (F(3,357)=4.44, p<0.01) and post hoc analysis revealed that MNM mice significantly decreased body weight on day 2 (p<0.05), day 3 (p<0.0001), and day 4 (p<0.0001) compared to baseline, and between MNM and SNM (day 2 p<0.05, day 3 p<0.05, day 4 p<001). Intriguingly, there was a day by drug interaction with female mice (F(2,297)=3.262, p<0.05), however, no changes in MNF mouse body weight were observed. Surprisingly, SNF mice demonstrated on average over a 1.5% increase in weight as compared to baseline (days 2-4 vs. baseline p<0.001), and post-hoc analysis revealed significant differences between SNF and MNF mice on days 2 and 4 (p<0.01, p<0.05 respectively, Fig. 1I). These results suggest that male and female mice both experience significant withdrawal under our paradigm, but that the behaviors and physiological changes experienced during this withdrawal are qualitatively different between the sexes.

### Morphine withdrawal increases BNST neuronal excitation in females, but not males

To understand how opioid withdrawal may alter the flow of information through the BNST, we performed whole-cell electrophysiology on putative BNST projection neurons (BNST_proj_, see methods) 24 hours after the last precipitated withdrawal. Examining the resting membrane potential (RMP), we found a drug by sex interaction between groups (F(1,79)=4.049, p<0.05; −41.59 ± 1.22 mV for SNM, n=19; −43.47 ± 1.48 mV for MNM, n=16, −48.19 ± 1.56 mV for SNF, n=26; −43.95 ± 1.51 mV for MNF, n=22, Fig. 2A). We next examined neuronal excitability in the BNST by measuring alterations in rheobase current. While we did not observe a drug by sex interaction here (p=0.22), we observed a main effect of morphine exposure (p<0.01, Fig. 2B). Planned t-tests showed that in males, morphine withdrawal did not alter the rheobase current (12 ± 1.58 pA for SNM, n=18; 9.4 ± 1.53 pA for MNM, n=15, p=0.25, Fig. 2B), however, it significantly decreased the rheobase current in female mice (15.15 ± 2.2 pA for SNF, n=23; 8.23 ± 0.8 pA for MNF, n=19, p<0.01, Fig. 2B). In concert, these results suggest that morphine withdrawal increases BNST_proj_ excitability in MNF, but not in MNM mice.

**Figure 2:**
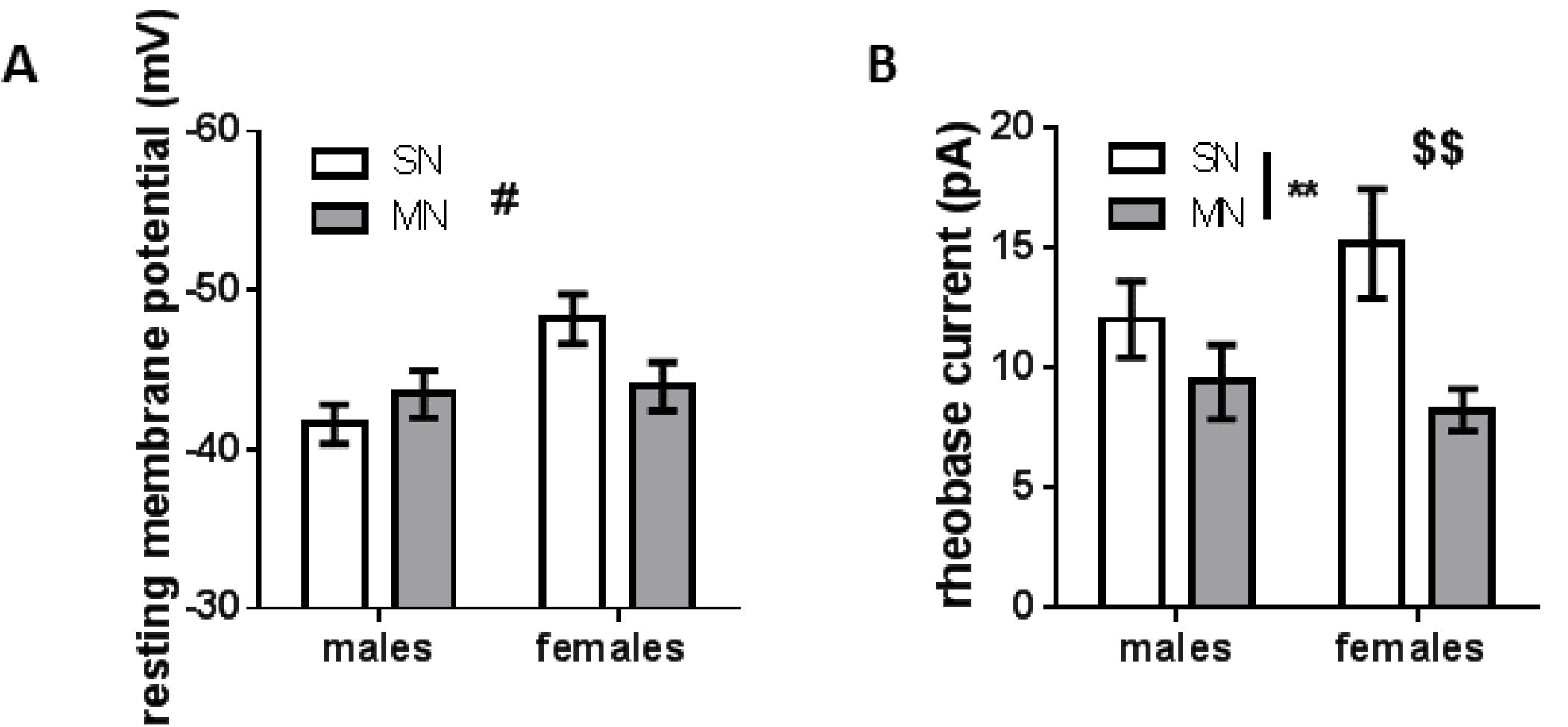
Morphine withdrawal alters properties of cell excitability in BNST_proj_. **A)** Bar graph of resting membrane potentials in BNST_proj_ (# drug by sex interaction p<0.05) **B)** Bar graph of rheobase current (main effect of drug treatment ** P<0.01; $$ planned t-test P<0.01, SNF n=23; MNF n=21). Each bar represents the mean ± SEM.

### Morphine withdrawal alters BNST GABAergic signaling in males and females

Due to the changes observed in excitability, and because within slice morphine withdrawal has been shown to enhance GABA transmission (Dumont and Williams, 2004), we hypothesized that there may be altered inhibitory tone driving these effects. Therefore, we investigated the ability of the *in vivo* withdrawal paradigm to induce a plastic change in synaptic inhibition on to BNST_proj_. In males, morphine withdrawal resulted in a significant leftward shift on the cumulative distribution curve for spontaneous-IPSC inter event interval (vs. SNM K-S test p<0.0001, Fig 3A). In contrast, morphine withdrawal in females resulted in a rightward shift on the cumulative distribution curve for spontaneous-IPSC inter event interval (vs. SNM K-S test p<0.0001, Fig. 3C). Furthermore, when we examined the within cell averages for frequency there was a significant drug by sex interaction (F(1,68)=10.26, p<0.01; 3.14 ± 0.51 Hz for SNM, n=14; 6.39 ± 1.06 Hz for MNM, n=22, p<0.05; 6.98 ± 0.93 Hz for SNF, n=17; 4.31 ± 0.77 Hz for MNF, n=19, p<0.05, Fig. 3E). In both males and females, we observed a trend for a decrease in sIPSC amplitude (cumulative distributions, K-S test, p=0.072) following morphine withdrawal (Fig. 3B and 3D). Examining cell averages, there was a main effect of sex (p<0.01), but no interaction (55.75 ± 8.14 pA for SNM n=14, 47.42 ± 5.45 pA for MNM, n=22; 37.88 ± 3.68 pA for SNF, n=17, 35.76 ± 3.50 pA for MNF, Fig. 3F.) These data suggest that repeated morphine withdrawal engages a bidirectional plasticity in GABAergic tone on BNST_proj_ in male and female mice, increasing GABA release in male mice and decreasing GABA release in female mice.

**Figure 3:**
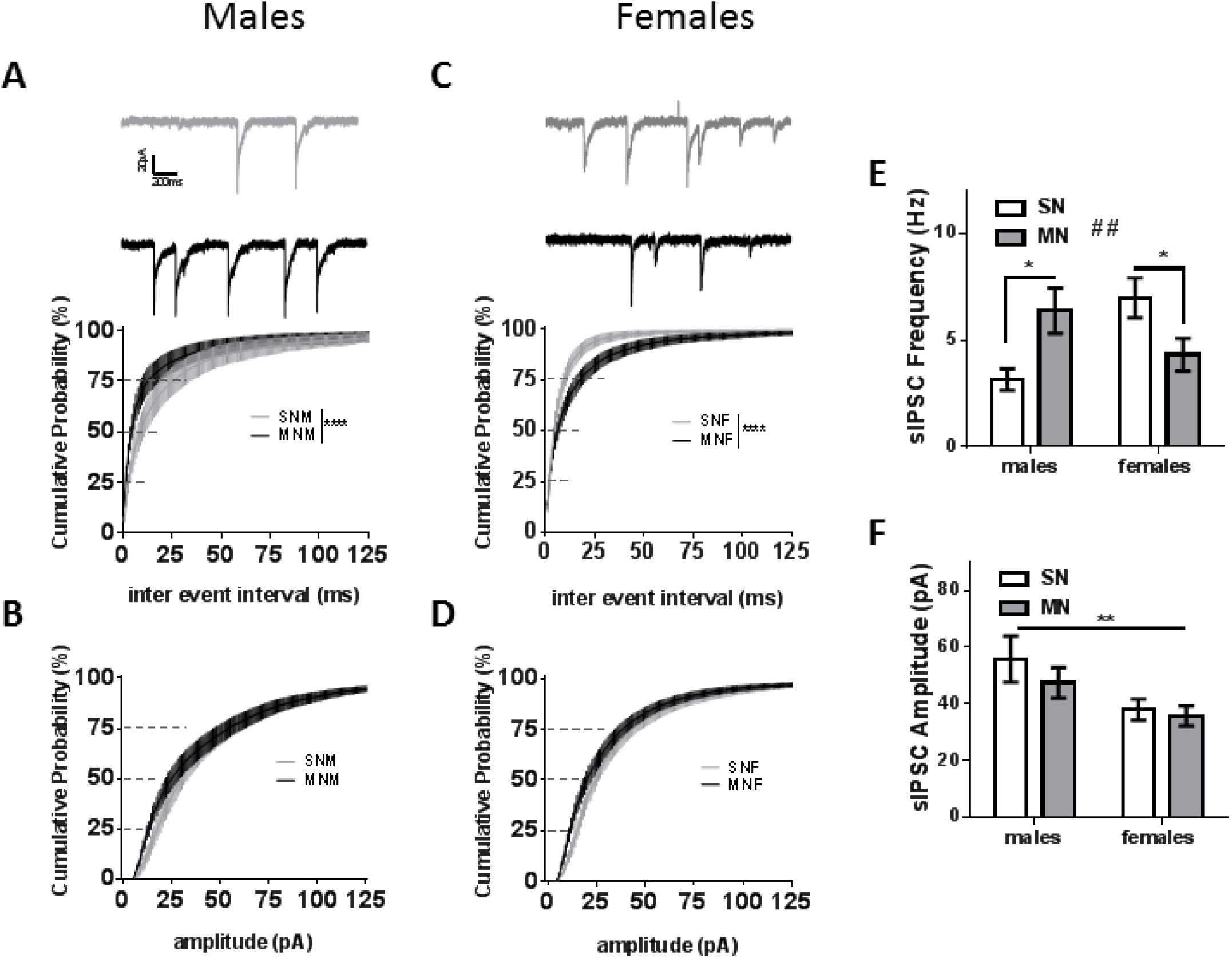
Morphine withdrawal differentially alters BNST spontaneous GABAergic transmission in male and female mice. **A)** Representative currents (SNM gray, MNM black) and cumulative distribution plot comparing sIPSC inter event interval in males (**** p<0.0001, SNM n=14; MNM n=22). **B)** Cumulative distribution plot comparing sIPSC amplitude in males from the same cells as A. **C)** Representative currents (SNF gray, MNF black) and cumulative distribution plot comparing sIPSC inter event interval in females (**** p<0.0001.SNF n=17; MNF n=19). **D)** Cumulative distribution plot comparing sIPSC amplitude in females from same cells as C. **E)** Bar graph of sIPSC frequency cell averages (## drug by sex interaction p<0.01, * P<0.05) **F)** Bar graph of sIPSC amplitude cell averages had main effect of sex (** p<0.01).

Previous research has shown that withdrawal can activate certain cell populations within the BNST (Dumont and Williams, 2004; Hamlin et al., 2004). To investigate the role of within slice excitability on morphine withdrawal induced plasticity of inhibitory transmission on BNST_proj_, we bath applied TTX (500 nM), to block action potentials and recorded mIPSCs 24 hours after the final naloxone injection. Similar to the sIPSC data, in males, we observed a leftward shift on the cumulative distribution curve for miniature-IPSC inter event interval (K-S test p<0.0001, Fig 4A). In contrast to sIPSCs, the effect of withdrawal on mIPSCs in female mice resulted in a leftward shift of the cumulative distribution curve (K-S test p<0.0001, Fig. 4C). Comparing within cell averages, we observed a strong trend for a main effect of sex (p=0.0825) but not a significant interaction (1.26 ± 0.22 Hz for SNM, n=19; 1.84 ± 0.25 Hz for MNM, n=12; 2.02 ± 0.58 Hz for SNF, n=13; 2.76 ± 0.68 Hz for MNF, n=15). Morphine withdrawal induced a rightward shift on the cumulative distribution curve for mIPSC amplitude in males, yet, similar to sIPSCs maintained a leftward shift in females (K-S test p<0.0001, Fig. 4B and 4D respectively). There was not a significant interaction, however, when within cell averages were examined (22.33 ± 2.20 pA for SNM, n=19; 24.79 ± 2.47 pA for MNM, n=12; 26.59 ± 1.85 pA for SNF, n=13; 26.54 ± 3.58 pA for MNF, n=15). The cumulative distributions suggest that the observed increase in the frequency of GABA release was apparent for both sexes following the addition of TTX and revealed opposite mIPSC amplitude profiles. These data suggest that within slice activity may partially account for the divergent excitability and sIPSC profile observed between BNST_proj_ in male and female mice following morphine withdrawal.

**Figure 4:**
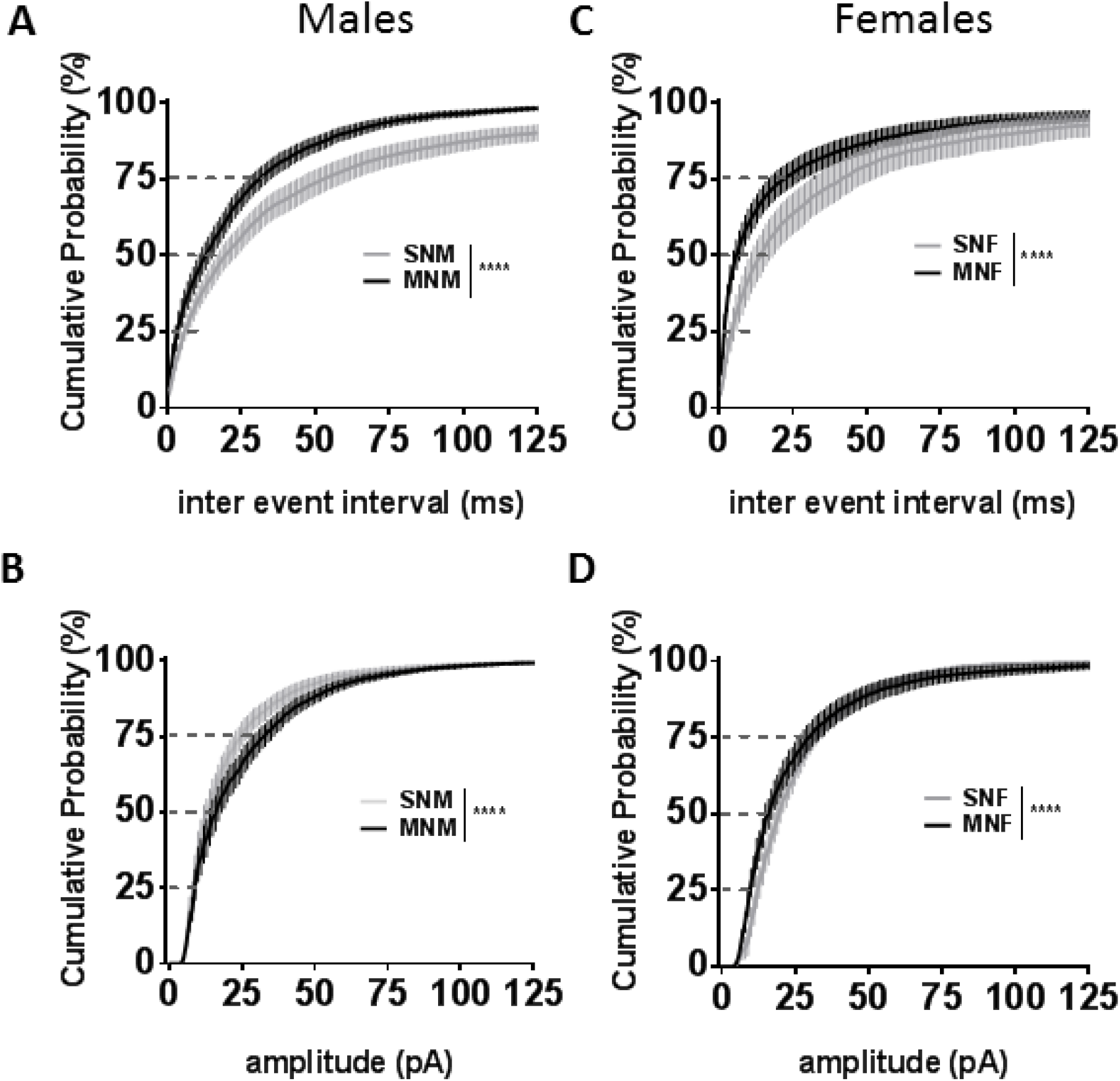
Alterations in BNST GABAergic signaling is unmasked by TTX in males and females. **A)** Cumulative distribution plot comparing miniature-IPSC inter event interval in male mice (SNM n=14; MNM n=22). **B)** Cumulative distribution plot comparing miniature-IPSC amplitude in male mice from the same cells as A. **C)** Cumulative distribution plot comparing miniature-IPSC inter event interval in female mice (SNF n=17; MNF n=19). **D)** Cumulative distribution plot comparing miniature-IPSC amplitude in female mice from the same cells as C. **** (p<0.0001).

To test this hypothesis, we drove within slice excitability directly using a paired-pulse electrical stimulation (50 ms inter-stimulus interval) and recorded evoked IPSCs. Here, we found a significant main effect of drug treatment (p<0.01). Furthermore, planned t-test showed morphine withdrawal significantly increased the paired-pulse ratio (PPR) in MNM compared to SNM (1.06 ± 0.08 PPR for SNM, n=12; 1.39 ± 0.1 PPR for MNM, n=20, p<0.05, Fig. 5C), and MNF compared to SNF (1.08 ± 0.07 PPR for SNF, n=22; 1.42 ± 0.12 PPR for MNF, n=19, p<0.05, Fig. 5C). These data suggest that promoting within slice activity reduces GABAergic synaptic efficacy in both sexes following withdrawal.

**Figure 5:**
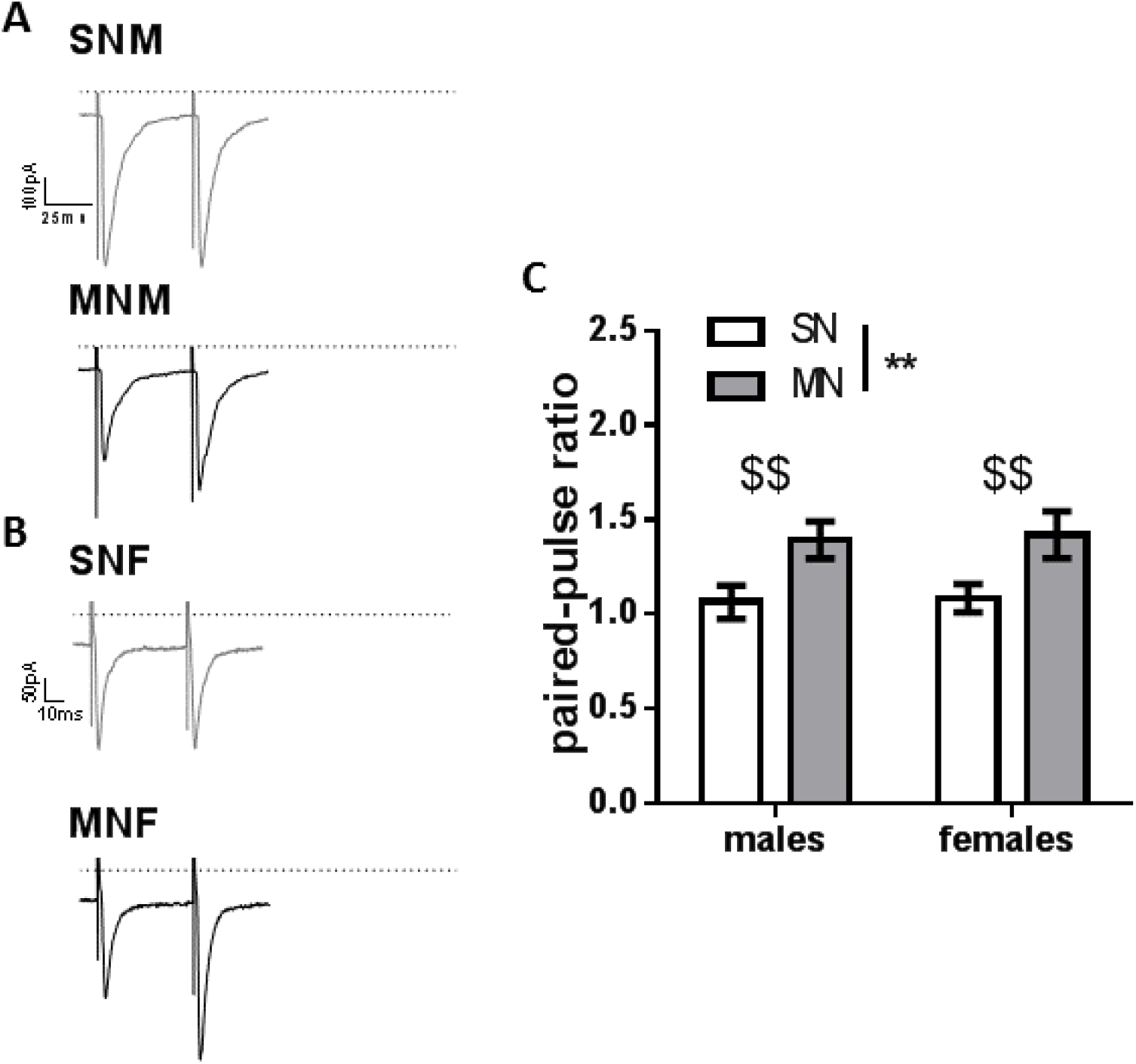
Morphine withdrawal increases the paired-pulse ratio in male and female mice. **A)** Representative paired-pulse traces from BNST neurons in male mice (SNM *top*; MNM *bottom*, dashed line 0 pA, scale bar 100 pA x 25 ms). **B)** Representative paired-pulse traces from BNST neurons in female mice (SNF, *top*; MNF *bottom*, scale bar 50 pA x 10 ms). **C)**. Bar graph of paired-pulse ratios (main effect of drug treatment, ** p<0.01; $$ planned t-tests p<0.01, SNM n=12; MNM n=20, SNF n=22; MNF n=19). Each bar represents the mean ± SEM.

## Discussion

In this study, we aimed to model repeated cycles of opioid withdrawal as a major contributor to the development of OUD (Koob and Volkow, 2016). Recent data suggests that 4 of 5 new heroin users previously used a prescription opioid (Cicero et al., 2014). Therefore, it is likely that the transition to an OUD occurs in patients who experience withdrawal from clinically prescribed opioids, and thus they may misuse opioids and potentially seek illicit avenues to continue usage. Because of this, we administered a dose of morphine known to be analgesic in male mice, 10 mg/kg s.c. (Dykstra et al., 2011) followed by naloxone (1 mg/kg) across 3 days to investigate adaptations. Although the contribution of sex on the analgesic potency of opioids has been examined previously (Kest et al., 1999), few studies have investigated withdrawal characteristics in male and female rodents (Blum et al., 1976; Cicero et al., 2002). Both MNM and MNF experienced significant somatic signs of withdrawal and robust sensitization of withdrawal across the 3-day paradigm, however, on average females exhibited more somatic symptoms (Fig. 1B & 1C). This was surprising, given the aforementioned decreased sensitivity to the analgesic properties of morphine observed in females and the relatively low dose of morphine administered. Additionally, weight loss has previously been described as a characteristic of morphine withdrawal (Goeldner et al., 2013; McElligott et al., 2013). Consistent with these studies, we found that MNM exhibited a 2.5% loss in body weight, however MNF did not lose weight across the paradigm (Fig. 1H and 1I). This decrease in body weight is likely not due to increased gut motility, because MNF produced significantly more fecal boli (Fig. 1E). This phenomenon in MNM may be attributed instead to a decrease in food consumption or an increase in energy expenditure. These data demonstrate that although there are differences in the analgesic properties of morphine between male and female mice, such analgesic differences do not necessarily correlate to experiences in withdrawal, suggesting that the physiological mechanisms in the brain underlying these behaviors may be sexually dimorphic.

A wealth of evidence suggests that the BNST acts as a central hub in the acute responses involved in opioid withdrawal, however, whether these changes lead to persistent alterations was unknown. Altering noradrenergic signaling within the BNST modulates both the somatic and affective components of morphine withdrawal (Delfs et al., 2000). Furthermore, the BNST is known to regulate feeding, fear, anxiety- and reward-like behaviors via projections to the VTA and LH (Jennings et al., 2013a,b; Marcinkiewcz et al., 2016). In male rodents, spontaneous and naloxone-precipitated withdrawal inhibits activity of VTA dopamine neurons (Diana et al., 1995; Georges et al., 2006) and hence, reduces dopamine release in the nucleus accumbens (Acquas et al., 1991; Kalivas, 1993). Additionally, studies implicate that orexin neurons in the LH contribute to somatic withdrawal behaviors (Georgescu et al., 2003; Sharf et al., 2008), and, inhibitory synaptic inputs from the BNST suppress the activity of LH glutamatergic neurons to regulate feeding (Jennings et al., 2013a). In the BNST, within slice morphine withdrawal increases GABAergic cell firing, enhancing the release of GABA. It was unknown however, if withdrawal could create plastic adaptations in inhibitory synapses that would persist 24 hours following the precipitation of withdrawal. Using a posthoc analysis, we focused our electrophysiology experiments on neurons putatively projecting to the VTA and LH exhibiting small capacitance, high input resistance, and lacking I_h_ current (see methods). In male mice, morphine withdrawal increased sIPSCs and mIPSCs frequency onto BNST_proj_ suggesting an action potential independent enhancement in presynaptic GABA release onto this circuit. Similarly, female mice experiencing morphine withdrawal had an enhancement of inhibition on BNST_proj_ when mIPSCs were examined; however, they exhibited decreased sIPSC and enhanced neuronal excitability in the absence of TTX. These data suggest that within slice excitability in the BNST of the female mice following morphine withdrawal drives the opposing s/mIPSC results. To test this hypothesis, we actively promoted excitability in the slice by electrical stimulation. While electrical stimulation releases the GABA that we actively measure, it will also potentially release a host of other signaling factors within the slice (Karkhanis et al., 2016; Stella et al., 1997). Using a paired pulse protocol with a 50 ms inter-stimulus interval, we found that morphine withdrawal increased the PPR of evoked GABAergic transmission onto the BNST_proj_ in both males and females. An increase in PPR is typically associated with a decrease in the release of neurotransmitter, similar to the observed decrease in sIPSC frequency in female mice. These data suggest that there are unknown factors within the slice that are released in an activity dependent fashion, which may regulate the GABAergic tone onto BNST_proj_. Furthermore, the data suggests that the release and/or abundance of these to-be-identified factors may be sexually dimorphic. Likely candidates for mediating this difference in GABAergic tone could be neuropeptides. Indeed, several neuropeptides investigated by our lab and others have shown differential effects on both pre- and postsynaptic GABAergic function (Kash and Winder, 2006; Krawczyk et al., 2011; Li et al., 2012; Normandeau et al., 2018; Pleil et al., 2015).

Finally, both male and female mice experiencing morphine withdrawal had a decrease in mIPSC amplitude on BNST_proj_, while also exhibiting an increase in mIPSC frequency. A decrease in mIPSC amplitude is typically consistent with a postsynaptic effect on GABA receptor function. The BNST contains a dense population of GABAergic interneurons (Kudo et al., 2012; Lebow and Chen, 2016) and receives GABAergic input largely through projections from the central nucleus of the amygdala (CeA) (Herman et al., 2013). Although we observe a host of changes in inhibition onto BNST_proj_, whether the observed alterations in GABAergic signaling stem from local BNST interneurons or through afferent input is unknown. Previous studies show that morphine withdrawal increases cFos expression, a marker of neuronal activation, in the BNST and CeA (Hamlin et al., 2004) and suggest that the CeA→BNST circuit contributes to the affective component of withdrawal (Nakagawa et al., 2005). Future studies will investigate the contributions of various GABAergic inputs onto BNST_proj_, and if the alterations we observe are occurring on disparate synapses.

Here we have presented evidence that both male and female mice exhibit significant withdrawal to a low dose of morphine, and that this withdrawal sensitizes over our 3-day paradigm. While female mice are less sensitive to the analgesic properties of morphine in general, they appear to have enhanced sensitivity to withdrawal as evidenced by increased somatic withdrawal symptoms, but a decreased sensitivity to the weight loss associated with morphine withdrawal. Repeated morphine withdrawal differentially regulates GABAergic tone onto BNST_proj_ in male and female mice, which may be due to within slice activity, and governs the excitability of BNST_proj_. Thus, our data highlight both behavioral and physiological sex differences to opioid withdrawal, and underscore the importance of conducting experiments in both sexes.

Authors’ contributions
B.R.L. conceived of project, performed behavior, performed electrophysiology, wrote manuscript; E.S.C. performed behavior, performed electrophysiology, edited manuscript; K.T.S. performed behavior, edited manuscript; D.P. performed electrophysiology, edited manuscript; M.M.P. performed electrophysiology, edited manuscript; K.D. performed behavior; Z.A.M. conceived of and supervised project, supervised BRL, ESC, KTS, KD, performed behavior, wrote manuscript

